# Significance of hydrophobic and charged sequence similarities in sodium-bile acid cotransporter and vitamin D-binding protein macrophage activating factor

**DOI:** 10.1101/2020.03.03.975524

**Authors:** Ibne Raihan Zunaid, Stefania Pacini, Marco Ruggiero

## Abstract

Sodium-bile acid cotransporter, also denominated sodium-taurocholate cotransporting polypeptide (NTCP) is an integral membrane protein with multiple hydrophobic transmembrane domains. The third extracellular domain of NTCP presents a stretch of nine aminoacids (KGIVISLVL) that is characterized by pronounced hydrophobicity and serves as receptor for a protein, preS1, showing the hydrophobic epta-peptide sequence NPLGFFP. Vitamin D-binding protein macrophage activating factor (DBP-MAF) is a multifunctional protein that is characterized by two hydrophobic regions able to bind fatty acids and vitamin D, respectively. Here we demonstrate that NTCP and DBP-MAF show significant sequence similarities as far as hydrophobic stretches of aminoacids are concerned. Alignment of the sequence of seven aminoacids preceding the 157-KGIVISLVL-165 stretch of NTCP shows four aminoacids that are identical to those of the corresponding sequence of DBP-MAF, and two that are conserved substitutions. In addition, in the sequence of DBP-MAF that is aligned with the sequence YKGIVISLVL of NTCP, there are two contiguous negatively charged aminoacids (ED) and, in the preceding epta-peptide sequence, there are three negatively charged aminoacids (D-ED), whereas in the corresponding sequence of NTCP there are only two (D--D) that are not contiguous. This concentration of negatively charged aminoacids may be involved in binding of protein inserts characterized by high density of positively charges residues. The alternating hydrophobic and electrostatic interactions described in this paper may help elucidating the biological roles of these proteins as far as protein-protein interactions are concerned.

## Introduction

The sodium-bile acid cotransporter, also defined the sodium-taurocholate cotransporting polypeptide (NTCP) or also liver bile acid transporter, is an integral transmembrane protein that is encoded by the SLC10A1 (solute carrier family 10, member 1) human gene and shows multiple hydrophobic transmembrane domains (1).

This transporter plays a critical role in maintenance of the enterohepatic recirculation of bile acids and hepatocyte function. The protein is constituted by 349 aminoacids with a resulting molecular mass of 56 kDa and works by binding two sodium ions and one (conjugated) bile salt molecule, in this manner determining hepatic influx of bile salts. Other molecules that may be transported by this protein comprise steroid and thyroid hormones, and xenobiotics of different origin (2). NTCP also represents the receptor for Hepatitis B virus (HBV) and the interaction between the aminoacid sequences of NTCP and those of HBV surface proteins has been characterized in molecular details. The preS1 sequence of large envelope protein (L) is essential for interaction with a stretch of nine hydrophobic residues (157-KGIVISLVL-165) located in the third transmembrane domain of NTCP. The sequence of the preS1 L protein binding NTCP is constituted by a stretch of predominantly hydrophobic aminoacids (residues 2-48) and, within this stretch, there is a highly conserved motif (9-NPLGF(F/L)P-15) that is essential for binding (3). It is apparent that knowledge of the mechanics of interaction at this level may be instrumental in developing strategies aimed at preventing preS1/NTCP interaction with consequent internalization. We therefore decided to use computational tools to investigate sequence similarities in other proteins that may interact with the preS1 sequence as well as with other surface proteins of viruses. To this end, we chose a well-characterized multifunctional serum protein that has the ability to work as a transporter, a hormone- and fatty acid binding protein, and an immune stimulating cytokine, the vitamin D-binding protein (DBP) macrophage activating factor (MAF). This is a serum alpha-2 glycoprotein constituted by a single polypeptide chain showing a molecular mass of 51–58 kDa. From the structural point of view, it is related to serum albumin and it is also known as Gc (Group-component) globulin. DBP synthesis occurs in the liver and its level in human plasma is around 20–55 mg/100 ml. DBP is a multifunctional protein that, in addition to vitamin D, binds actin, acting as an actin scavenger, and also binds fatty acids. The protein is constituted by 458 aminoacids and shows three domains that share limited sequence similarity with each other as well as with similar sequences of human albumin. The first domain contains the vitamin D-binding site that is located in a shallow cleft that allows interaction with the plasma membrane. DBP is present on the surface of several cell types that comprise yolk sac endodermal cells and T lymphocytes. In B lymphocytes, DBP is responsible for the linkage of surface immunoglobulins, thus contributing to the balance of the immune response (4). The de-glycosylated form of DBP is a powerful macrophage activating factor - hence the designation DBP-MAF - and shows a number of biological effects (5). We demonstrated that, in addition to its presence in human serum, DBP-MAF is formed during fermentation of milk and colostrum (6) as well as during fermentation of hemp seed proteins (7). Here, we demonstrate that DBP-MAF shares significant similarities with hydrophobic sequences of NTCP that interact with the preS1 sequence of L protein. We also show that DBP-MAF, but not NTCP, shares similarities between the hydrophobic sequence of the fatty acid binding site and the domain IV of the same L protein. Finally, we show the differences in the distribution of negatively charged aminoacids in DBP-MAF and NTCP and we discuss their biological implications.

## Materials and Methods

Study of sequence similarities were performed using the Align tool of Uniprot (uniprot.org). For NTCP, the sequence UniProtKB - Q14973 (NTCP_HUMAN) was used. For DBP-MAF, the sequence UniProtKB - P02774 (VTDB_HUMAN) was used. For HBSAG, the sequence UniProtKB - P03138 (HBSAG_HBVD3. Isolate France/Tiollais/1979) was used. For alpha-N-acetylgalactosaminidase (abbreviated in NAGAB_HUMAN), the sequence UniProtKB - P17050 (NAGAB_HUMAN) was used. The sequences TNGTKR, HKNNKS, RSYLTPGDSSSG, and QTNSPRRA were retrieved from Jaimes et al. (10). The yellow color used in Fig 1 indicates the transmembrane domains of NTCP. The gray color indicates similarities; the conventional consensus symbols are: "*" indicating that the residues are identical in all sequences in the alignment. ":" indicating that conserved substitutions have been observed. "." indicating that semi-conserved substitutions are observed, that is, amino acids having similar features. The light indigo color indicates hydrophobicity. The pale red color indicates negatively charged residues. Study of *in vitro* interaction between DBP-MAF or the product of microbial fermentation of milk and colostrum mentioned above, here abbreviated in PMF (Product of Microbial Fermentation), and NAGAB_HUMAN was was performed by R.E.D. Laboratories (Zellik, Belgium) where NAGAB_HUMAN and DBP-MAF had been purified. NAGAB_HUMAN is an enzyme that cleaves terminal alpha-N-acetylgalactosamine residues from glycolipids, glycopeptides and glycoproteins; it binds to the alpha-N-acetylgalactosamine moiety that is attached to threonine in the sequence TPTELAK close to the carboxyl terminus of DBP-MAF. Therefore, binding of NAGAB_HUMAN is a direct method to evaluate DBP-MAF activity in serum or other matrices. For these experiments, microtiter plates coated with a specific antibody directed against NAGAB_HUMAN were used. Samples were incubated with standardized dilutions of NAGAB_HUMAN coming from a pool of 300 human sera from healthy subjects who did not show any sign of chronic inflammation or any disease. After 1 h incubation and exhaustive washing, complexes formed by NAGAB_HUMAN and purified DBP-MAF, used as positive control, or NAGAB_HUMAN and PMF, were revealed with horse radish peroxidase conjugate to rabbit antibody. The serum pool was mixed either with 200 ng of purified DBP-MAF, or with the fermented product in phosphate buffered saline (PBS) with the goal of studying the kinetics of interaction between NAGAB_HUMAN and DBP-MAF or NAGAB_HUMAN and the PMF. The compounds were incubated at 25C for 4, 24, 48, 72 and 120 h. Values for NAGAB_HUMAN binding activity in the absence of DBP-MAF or the PMF, with only PBS in the reaction mixture, were taken as 1.00. The experiment was repeated twice and the results reported in this study are the means of the two experiments.

**Figure 1.**
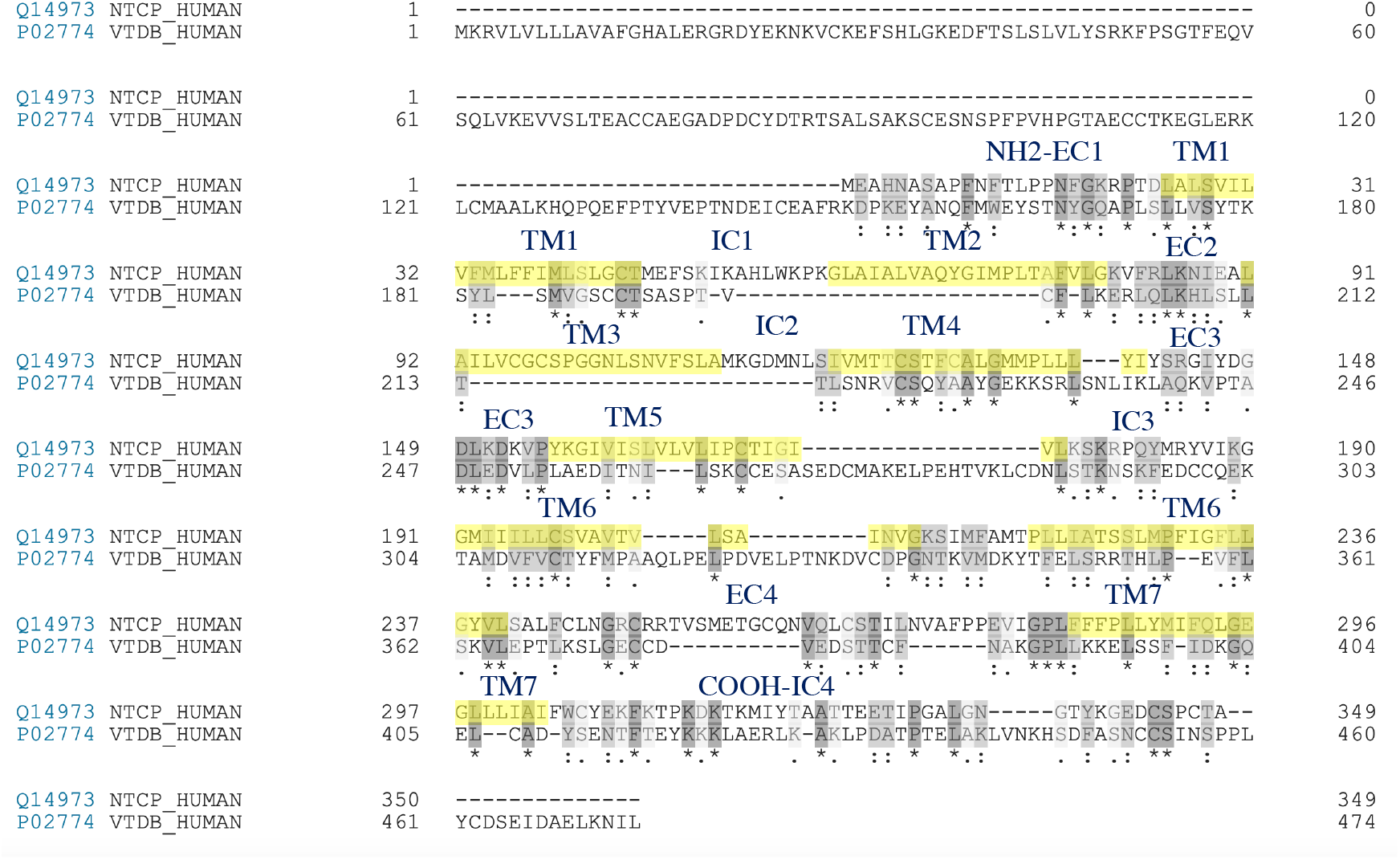
Sequence alignment between sodium-taurocholate cotransporting polypeptide (NTCP) and vitamin D-binding protein macrophage activating factor DBP-MAF. The sequences of NTCP (Q14973 NTCP_HUMAN) and DBP-MAF (also known as Gc-protein-derived macrophage activating factor; P02774 VTDB_HUMAN) were obtained and aligned using the Align tool of Uniprot (uniprot.org). The seven transmembrane domains of NTCP are highlighted in yellow and numbered TM1-TM7. The four extracellular domains starting from the amino terminus are numbered EC1-EC4. Likewise, the four intracellular domains are numbered IC1-IC4. The gray color indicates similarities. The conventional consensus symbols are: "*" indicating that the residues are identical in all sequences in the alignment. ":" indicating that conserved substitutions have been observed. "." indicating that semi-conserved substitutions are observed, that is, aminoacids having similar features.

## Results and Discussion

Fig. 1, shows the results of alignment of NTCP and DBP-MAF. The seven transmembrane domains of NTCP are highlighted in yellow and numbered TM1-TM7. The four extracellular domains starting from the amino terminus are numbered EC1-EC4. Likewise, the four intracellular domains are numbered IC1-IC4, with the last domain pertaining to the intracellular carboxyl terminus. The degree of similarity in the aminoacid sequences is apparent in several regions of the proteins.

Fig. 2, panel A, shows the infectivity determinant of HBV. The preS1 sequence in L protein (residues 2-48) is the sequence that specifically interacts with NTCP. Within this sequence, a highly conserved motif (9-NPLGF(F/L)P-15) is crucial for binding (11). Fig. 2, panel B, shows the preS1 binding sequence of NTCP aligned with the corresponding sequence of DBP-MAF. It is worth noticing that the sequence of seven aminoacids preceding the 157-KGIVISLVL-165 stretch of NTCP, that is the stretch that binds preS1, shows four amino acids that are identical to those of the aligned sequence of DBP-MAF, and two that are conserved substitutions. Fig. 2, panel C, shows that DBP-MAF presents a sequence that is very similar to that of domain IV of the L protein of HBV. Both sequences are rich in hydrophobic residues; the sequence of DBP-MAF is located in a shallow cleft of the proteins that facilitates its binding to fatty acids that are sandwiched between the protein and the plasma membrane as we proposed in 2013 (12). The sequence of domain IV of the L protein of HBV is a transmembrane domain that participates in infectivity (11). Fig. 2, panel D, shows the peculiar concentration of negatively charged residues in the sequence of DBP-MAF that is aligned with the sequence of NTCP that binds preS1. In DBP-MAF, there are two contiguous negatively charged amino acids (ED) and, in the preceding epta-peptide sequence, there are three negatively charged aminoacids (D-ED) for a total of five negatively charged residues, whereas in the corresponding sequence of NTCP there are only two (D--D) that are not contiguous. This concentration of negatively charged aminoacids may be involved in the binding and electrostatic neutralization of protein inserts characterized by high density of positively charges residues. In particular, it may be involved in binding the sequences TNGTKR, HKNNKS, RSYLTPGDSSSG, and QTNSPRRA that are characterized by a high surface concentration of positive charges interspersed with hydrophobic residues (10, 13). It is worth noticing that neutralization of positively charged residues may represent a strategy that exploits electrostatic interactions between charges on the surface of molecules. Such as strategy may not be limited to proteins; negatively charged glycosaminoglycans such as low-molecular-weight chrondroitin sulfate (14) might also serve the scope of binding and neutralizing those sequences.

**Figure 2.**
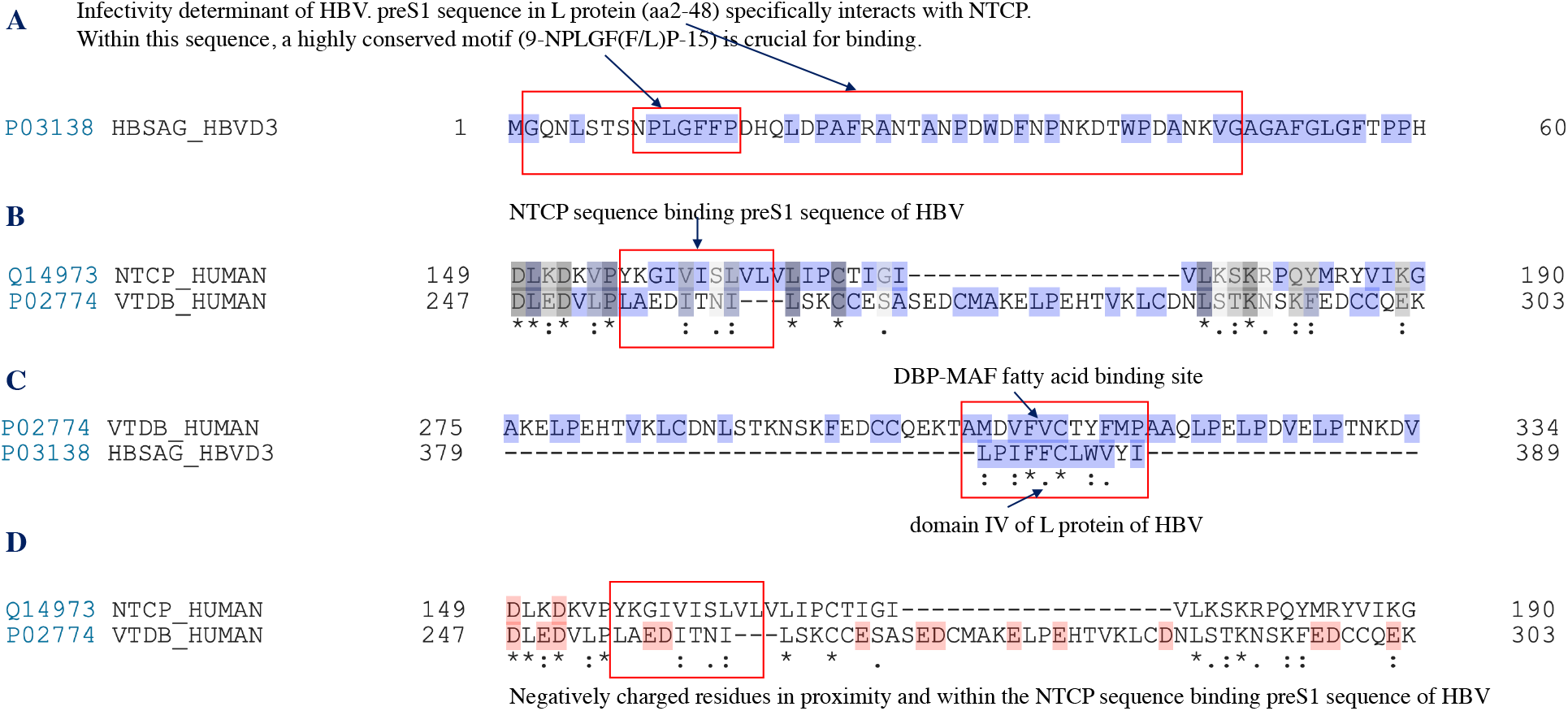
Sequence alignment between specific sequences of sodium-taurocholate cotransporting polypeptide (NTCP), vitamin D-binding protein macrophage activating factor DBP-MAF, and HBSAG. For HBSAG, the sequence UniProtKB-P03138 (HBSAG_HBVD3. Isolate France/Tiollais/1979) was used. The light indigo color indicates hydrophobicity. The pale red color indicates negatively charged residues.

Fig. 3, shows the results of alignment of NTCP, DBP-MAF and NAGAB_HUMAN. This latter protein has enzymatic activity and directly interacts with DBP-MAF, more precisely, it binds to alpha-N-acetylgalactosamine that is attached to threonine in the sequence TPTELAK close to the carboxyl terminus of DBP-MAF (see the sequence within the red square in Fig. 3). We chose to study this protein because of its role in DBP-MAF function and we found that there is a high degree of similarity with the other two proteins, DBP-MAF and NTCP, with particular reference to stretches of hydrophobic aminoacids. Based on these evidence, we asked an independent laboratory to perform *in vitro* experiments to assess the binding of DBP-MAF to NAGAB_HUMAN using, as ligands, purified DBP-MAF and PMF (6,8).

**Figure 3.**
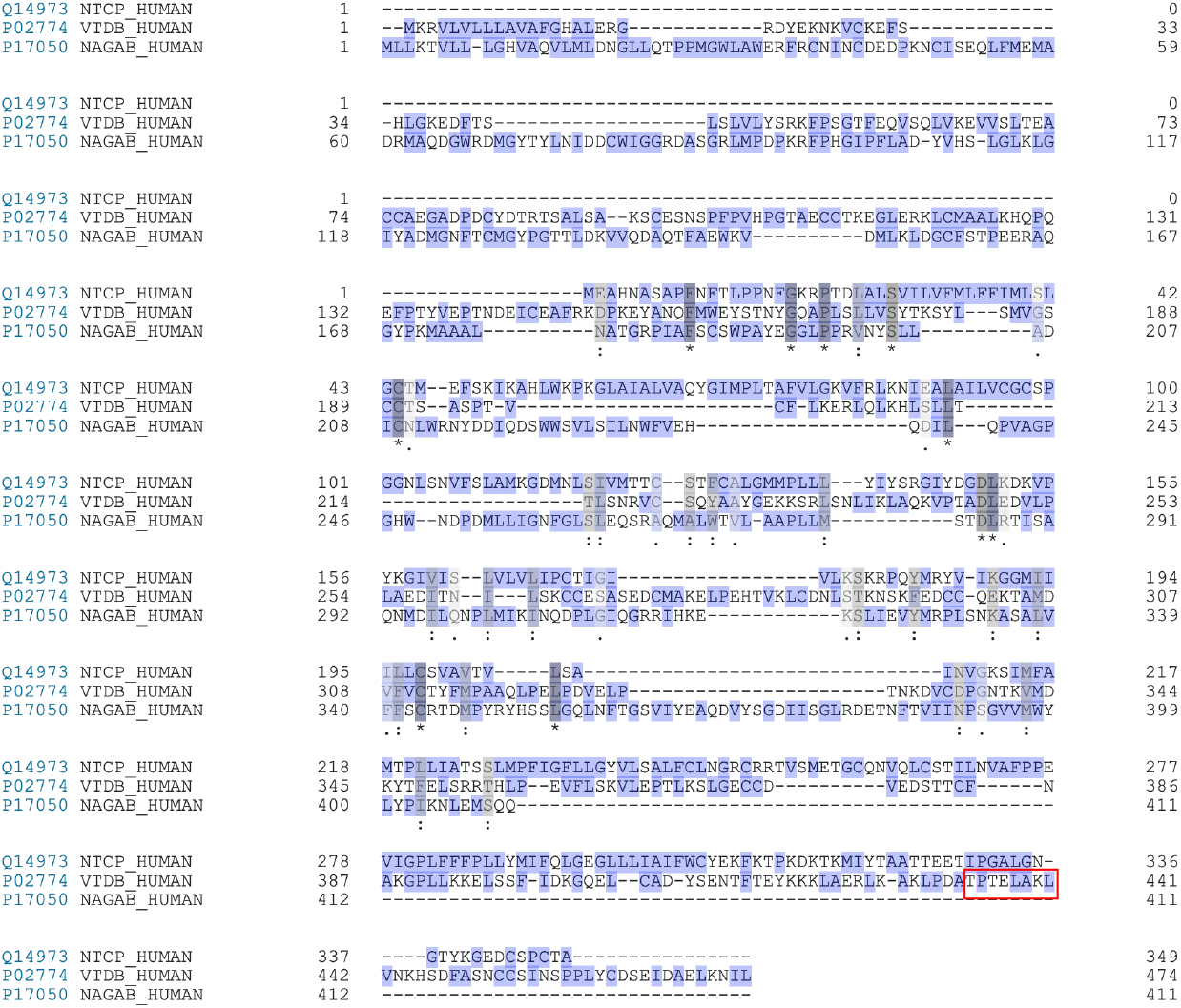
Sequence alignment between specific sequences of sodium-taurocholate cotransporting polypeptide (NTCP), vitamin D-binding protein macrophage activating factor DBP-MAF, and human alpha-N-acetylgalactosaminidase (NAGAB_HUMAN). For the latter, the sequence UniProtKB-P17050 (NAGAB_HUMAN) was used.

Fig. 4, shows that purified DBP-MAF, used as positive control, bound NAGAB_HUMAN only after 4 h incubation, reached a peak at 48 h, and returned below baseline values at 120 h. PMF, diluted 1:10 with PBS, had an initial value (at time 0) higher than that observed with PBS alone and higher than that observed with purified DBP-MAF; we interpret this result as indication that PMF has a much higher NAGAB_HUMAN-binding activity in comparison with purified DBP-MAF. At every time point, the PMF showed significantly higher activity in comparison with purified DBP-MAF. At 120 h, the activity of PMF was still well above the value obtained with the negative control, PBS, whereas DBP-MAF did not show any residual activity (not shown). Other experiments performed with 1:100 dilution showed a similar trend and the activity of PMF was always higher than that of purified DBP-MAF even at the highest dilution. These results led us to conclude that PMF has a DBP-MAF activity more than 100 fold higher than that of purified DBP-MAF. We attribute this higher activity to the presence of vitamin D_3_ and fatty acids that are natural constituents of milk and colostrum. The presence of these hydrophobic moieties may favor interaction between DBP-MAF and NAGAB_HUMAN stabilizing the complex formed by the two proteins.

**Figure 4.**
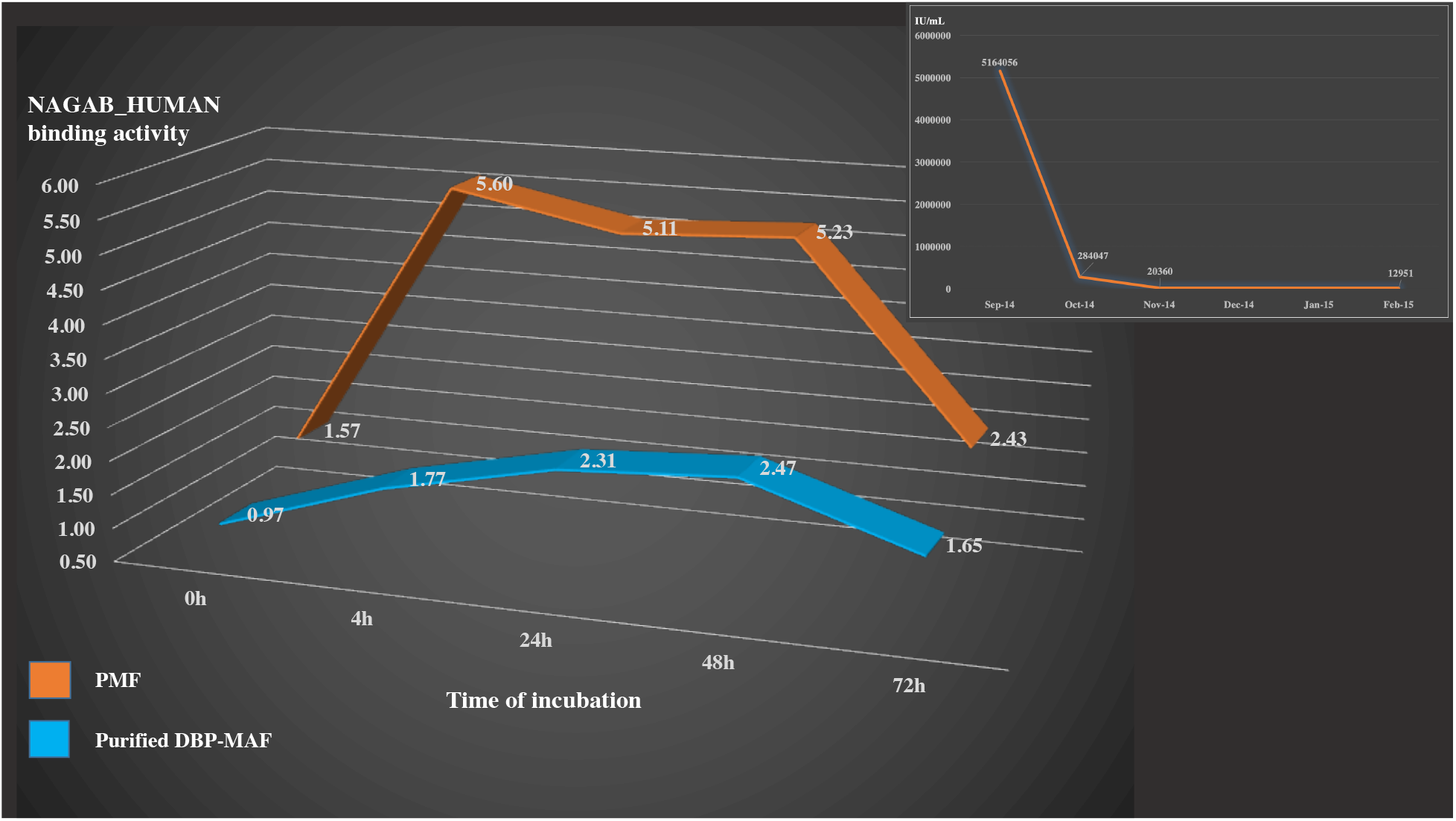
Kinetics of binding of NAGAB_HUMAN and purified DBP-MAF or product of microbial fermentation (PMF). Values for NAGAB_HUMAN binding activity in the absence of DBP-MAF or the PMF, with only PBS in the reaction mixture, were taken as 1.00. Insert: decrease of HBV viral load (IU/mL) after assumption of PMF. Sep-14 indicates values before assumption. Data obtained by three independent laboratories.

The sequence similarities shown in Fig. 3 seem to support this hypothesis that is further corroborated by the recent observation that DBP-MAF conjugated with vitamin D_3_ is much more efficient than DBP-MAF alone as an immune stimulating factor (15). Hydrophobic interactions between the sequences of DBP-MAF and preS1 and domain IV of L protein may also help interpreting the biological effects reported in Fig. 4 (insert). In this case, we can hypothesize two binding sites located at the two extremities of L protein. These interactions would be stabilized, as in the case of DBP-MAF/NAGAB_HUMAN complexes, by vitamin D_3_ and fatty acids. This hypothesis is corroborated by the observation that viral proteins are able to bind directly vitamin D_3_ (16), and fatty acids show anti-viral properties that are due to prevention of entry of the viral genome into the host cell (17) as we propose being the case for the interaction between DBP-MAF/vitamin D_3_/fatty acids and preS1 and L protein hydrophobic sequences. In conclusion, based on the *in silico* and *in vitro* observation reported in this study, we propose that vitamin D_3_ and fatty acids play a pivotal role in facilitating hydrophobic interactions between proteins showing stretches of similarities.

## Acknowledgements

The Authors wish to thank Dr. Tanja Mijatovic, PhD, Chief Scientific Officer at R.E.D. Laboratories, for insights on the significance of laboratory tests and for contributing to the understanding of the role of NAGAB_HUMAN in health and disease.

## Advisory

Any medical or scientific information present in this paper is provided for research, educational and informational purposes only. It is not in any way intended or implied to be used as a substitute for professional medical advice, diagnosis, treatment or care of any disease mentioned or implied in this study.

